# Synthetic lethality in large-scale integrated metabolic and regulatory network models of human cells

**DOI:** 10.1101/2023.01.27.525829

**Authors:** Naroa Barrena, Luis V. Valcárcel, Danel Olaverri-Mendizabal, Iñigo Apaolaza, Francisco J. Planes

## Abstract

Synthetic lethality (SL) is a promising concept in cancer research. A wide array of computational tools has been developed to predict and exploit synthetic lethality for the identification of tumour-specific vulnerabilities. Previously, we introduced the concept of genetic Minimal Cut Sets (gMCSs), a theoretical approach to SL for genome-scale metabolic networks. The major challenge in our gMCS framework is to go beyond metabolic networks and extend existing algorithms to more complex protein-protein interactions. We present here a novel computation approach that adapts our previous gMCS formulation to incorporate linear regulatory pathways. Our novel approach is applied to calculate gMCSs in integrated metabolic and regulatory models of human cells. In particular, we integrate the most recent genome-scale metabolic network, Human1, with 3 different regulatory network databases: Omnipath, Dorothea and TRRUST. Based on the computed gMCSs and transcriptomic data, we detail new essential genes and their associated synthetic lethals for different cancer cell lines. The performance of the different integrated models is assessed with available large-scale *in-vitro* gene silencing data. Finally, we discuss the most relevant gene essentiality predictions based on published literature in cancer research.

## Introduction

Two (or more) genes are synthetic lethal when the loss of function of either gene on its own is compatible with cell viability, while the co-occurrence of them leads to cellular death (O’Neil *et al.*, 2017). Given the plethora of tumour-specific genetic alterations, synthetic lethality (SL) is an attractive approach to identify selective and novel drug targets in cancer cells. This has propelled the development of robust methods to identify synthetic lethals from very different perspectives (Jerby-Arnon et al., 2014; Blomen et al., 2015; Lee et al., 2018; Zhang et al., 2021; Gimeno et al., 2022).

In previous works (Apaolaza et al., 2017; Apaolaza et al. 2019), we introduced the concept of genetic Minimal Cut Sets (gMCSs), a theoretical approach to SL based on genome-scale metabolic networks. gMCSs define minimal set of gene knockouts that blocks a particular metabolic task, typically the biomass reaction in cancer studies. They can be easily integrated with -omics data and used to elucidate metabolic vulnerabilities in cancer cells. Recently, based on data from the Cancer Dependency Map (DepMap) (Ghandi *et al.*, 2019; Meyers *et al.*, 2017), we assessed the capacity of our gMCS approach to predict gene essentiality in cancer cell lines and reported a superior performance than other network-based algorithms (Valcárcel *et al.*,2022). In a different work (Apaolaza *et al.*, 2022), we also integrated nutritional perturbations into our gMCS framework, leading to nutrient dependencies in cancer cell lines.

Unfortunately, our current gMCS framework is constrained to the metabolic space, which represents only a fraction of all the interactions that occur within a cell. For instance, the latest reconstruction of the human metabolism, Human 1 (Robinson *et al.*, 2020), only represents 22% of the genes available in Omnipath (Türei *et al.*, 2016), one of the biggest protein-protein interactions database. For this reason, the main challenge of our gMCS approach is to go beyond metabolic networks and extend existing algorithms to more complex protein-protein interactions, such as signalling or regulatory networks.

Our gMCS approach is built on gene-protein-reaction (GPR) rules available in genome-scale metabolic models (Ponce-De-León *et al.*, 2020). A natural way to extend our gMCS formulation is to incorporate regulatory information into these GPR rules, as done in other constraint-based modelling tools (Chandrasekaran and Price, 2010; Marmiesse et al., 2015; Wang et al., 2017). However, GPR rules in metabolic models are simple Boolean networks without negation terms and cycles, which are typically present in regulatory networks. This fundamental difference makes particularly challenging the integration of regulatory networks with our gMCS approach, which currently cannot deal with Boolean equations involving negation terms and cycles (Apaolaza *et al.*, 2019).

Here, we present a novel computation approach that adapts our previous gMCS formulation to incorporate linear regulatory pathways. Our novel approach is applied to calculate gMCSs in integrated metabolic and regulatory models of human cells. In particular, we consider Human1 (Robinson *et al.*, 2020) with 3 different regulatory network databases: Omnipath (Türei *et al.*,2016), Dorothea (Garcia-Alonso *et al.*, 2019) and TRRUST (Han *et al.*, 2018). Based on the computed gMCSs and transcriptomic data, we detail new essential genes and their associated synthetic lethals for different cancer cell lines. The performance of the different integrated models is assessed with available large-scale *in-vitro* gene silencing data (Hart *et al.*, 2015; Ghandi *et al.*, 2019; Meyers *et al.*, 2017). Finally, we discuss the most relevant gene essentiality predictions based on published literature in cancer research.

## Methods

### Enumeration of gMCSs via Mixed-Integer Linear Programming

Assume we have a metabolic network involving *m* metabolites and *n* reactions. This is typically represented with the stoichiometry matrix *S*, where each column represents a different reaction and each row a single metabolite. Reaction products and substrates have positive and negative coefficients, respectively. The flux vector *r* denotes the activity of the reactions. Here, reversible reactions were split into two irreversible steps and, therefore, reaction fluxes are non-negative (Eq. (1)).

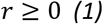

The application of the mass balance equation under steady state leads to Eq. (2), where the sum of fluxes which produce a certain metabolite is equal to the sum of fluxes which consume it.

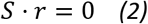

Our objective is to block a given metabolic task making use of the least number of gene knockouts. The metabolic task to disrupt can be represented as in Eq. (3):

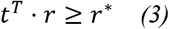

being *t* a null vector with a 1 in the position of the reactions involved in the metabolic task to target and *r*\* a positive constant.

In order to calculate gMCSs, *i.e.* minimal subsets of gene knockouts that disrupt an essential metabolic task, we need to define the possible gene knockout constraints, which take the following form:

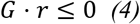

where the binary *G* matrix, of dimensions *l*x*n*, defines for each row *i* the set of blocked reactions, *G(i)*=*{k|G_ik_*=*1}*, arising from the knockout of an irreducible subset of genes. The subset of genes associated with each row in *G* is interrelated and their simultaneous knockout is required to delete at least one of the reactions in the metabolic network. This information is stored in the binary matrix *F* of dimensions *l*x*g*, which defines the subset of gene deletions involved in each row *i* in *G*, *F(i)*=*{p|F_ip_*=*1}*. In other words, the deletion of genes in *F(i)* leads to the disruption of reactions in *G(i).* An example metabolic network, including gene-protein-reaction (GPR) rules, can be found in Figure 1a. For illustration, Figure 1b show its associated *G* and *F* matrices, where, according to their second row, the knockout of gene 2 (*g_2_*) leads to the blockage of reaction 2 and 3 (*r*_2_, *r*_3_).

**Figure 1.**
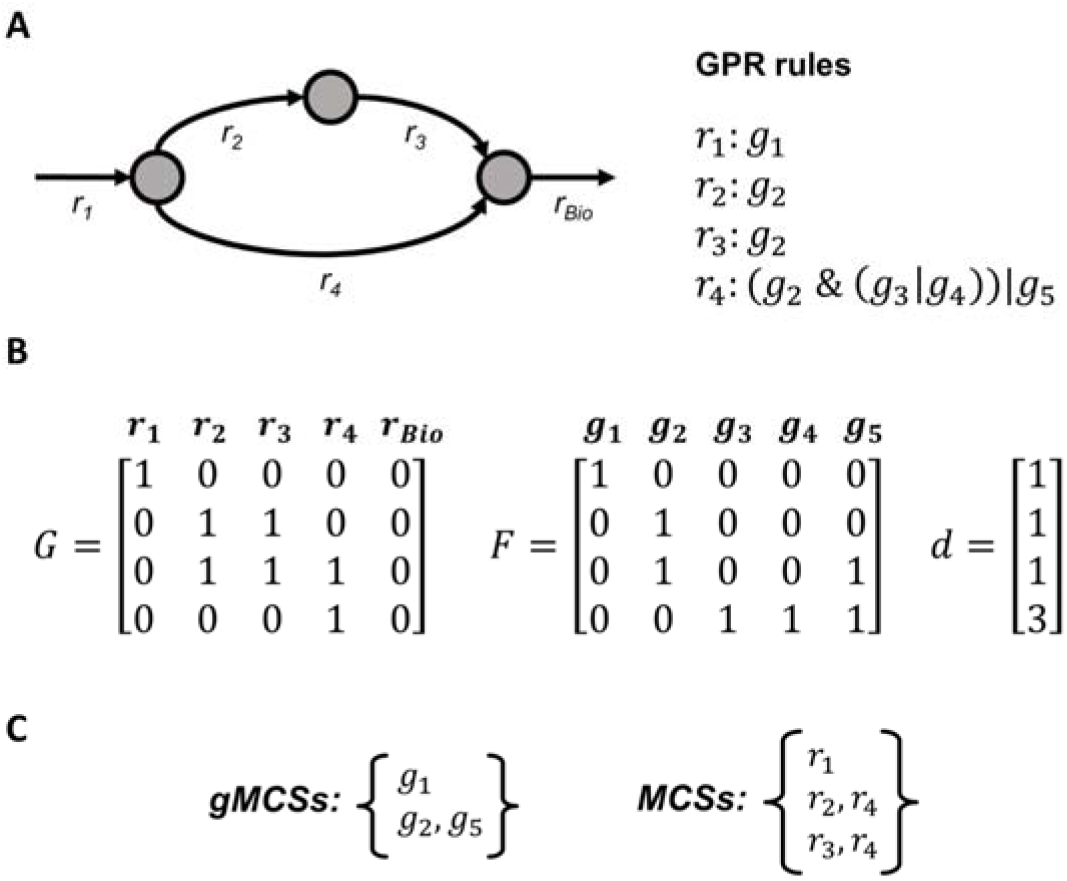
Illustration of gMCSs and MCSs. a) Example metabolic network and GPR rules. It involves 5 reactions (*r*_1_, *r*_2_, *r*_3_, *r*_4_, *r*_bio_) and 5 genes (*g*_1_, *g*_2_, *g*_3_, *g*_4_, *g*_5_); b) Matrices of gene knockout constraints, *G* and *F*, and net contribution of each row in *G* in terms of gene knockouts, *d.* For example, the second row in *G* is dependent on the third row in *G* and, thus, *d_3_*=2-1=1; c) Resulting set of gMCSs and MCSs.

From the infeasible primal problem defined by Eqs. (1)-(4), we formulate the unbounded dual problem and minimize the number of gene knockouts to block the target metabolic task with the following mathematical model:

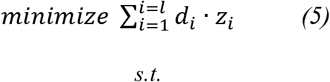

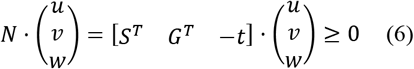

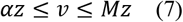

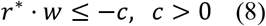

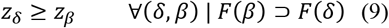

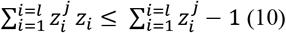

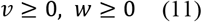

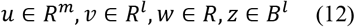

where *u, v* and *w* are dual variables associated with the mass balance equation, gene knockout constraints and the target metabolic task equation, respectively; *z* are binary variables linked to *v* through Eq. (7), namely *z*=0 ↔ *v*=0, *z*=1 ↔ *v*>0. Note here that *α* and *M* are small and large positive constants, respectively. Eq. (8) forces *w* to be non-zero, which makes the target metabolic task equation part of the infeasible primal problem. Eq. (9) considers the dependencies between dual variables *v* that may lead to non-minimal solutions, as it is described in Apaolaza et al., 2019. In addition, *d* is a known vector storing the number of gene deletions exclusively provided by its associated dual variable *v* and not by its dependent dual variables (see Figure 1b for illustration). Dependencies between dual variables can be easily obtained from *F* matrix. Finally, Eq. (10) allows us to eliminate previously obtained solutions (*z^j^*) from the solution space and identify new gMCSs.

In summary, the mixed-integer linear program defined by Eqs. (5)–(12) (MILP1) allows us to enumerate gMCSs in increasing order of gene knockouts. Figure 1c shows the resulting set of gMCSs for the example network considered. Note here that a similar approach can be built for Minimal Cut Sets (MCSs), which involves reaction knockouts instead of gene knockouts, as developed in different works (von Kamp and Klamt, 2014) (Figure 1c). In particular, for the computation of MCSs, the matrix *G* in Eq. (6) becomes the identity matrix (if all reactions are irreversible) and, thus, dependency constraints in Eq. (9) can be neglected.

### Calculation of G matrix in metabolic networks

MILP1 requires as input data different matrices: *S, G, F* and *t.* The construction of *G* and *F* matrices is not a trivial task, as demonstrated in Apaolaza et al., 2019 where we presented an efficient algorithm for their computation in complex metabolic networks. This technical improvement has allowed us to enumerate thousands of gMCSs in genome-scale metabolic networks in human cells (Valcárcel *et al.*, 2022).

Our *G* matrix construction algorithm involves 2 stages: i) calculation of irreducible subsets of gene knockouts that block each reaction separately using GPR rules; ii) integration of these irreducible subsets for the definition of *G* and *F* matrices. The first stage is the most challenging part, but it could be elegantly solved by transforming GPR rules into artificial reaction networks, called here GPR networks, and apply the MCS approach to block the target reaction(Apaolaza *et al.*, 2019) only considering the deletion of exchange reactions. Figure 2a shows the GPR rule for reaction 4 (*r_4_*) present in the example in Figure 1, the associated GPR network and the 2 resulting MCSs. This strategy could be followed because GPR rules define Boolean networks that do not involve **i)** negation (inhibition) terms and **ii)** cycles that could lead to oscillatory behaviour, as it is typically found in complex regulatory networks.

**Figure 2.**
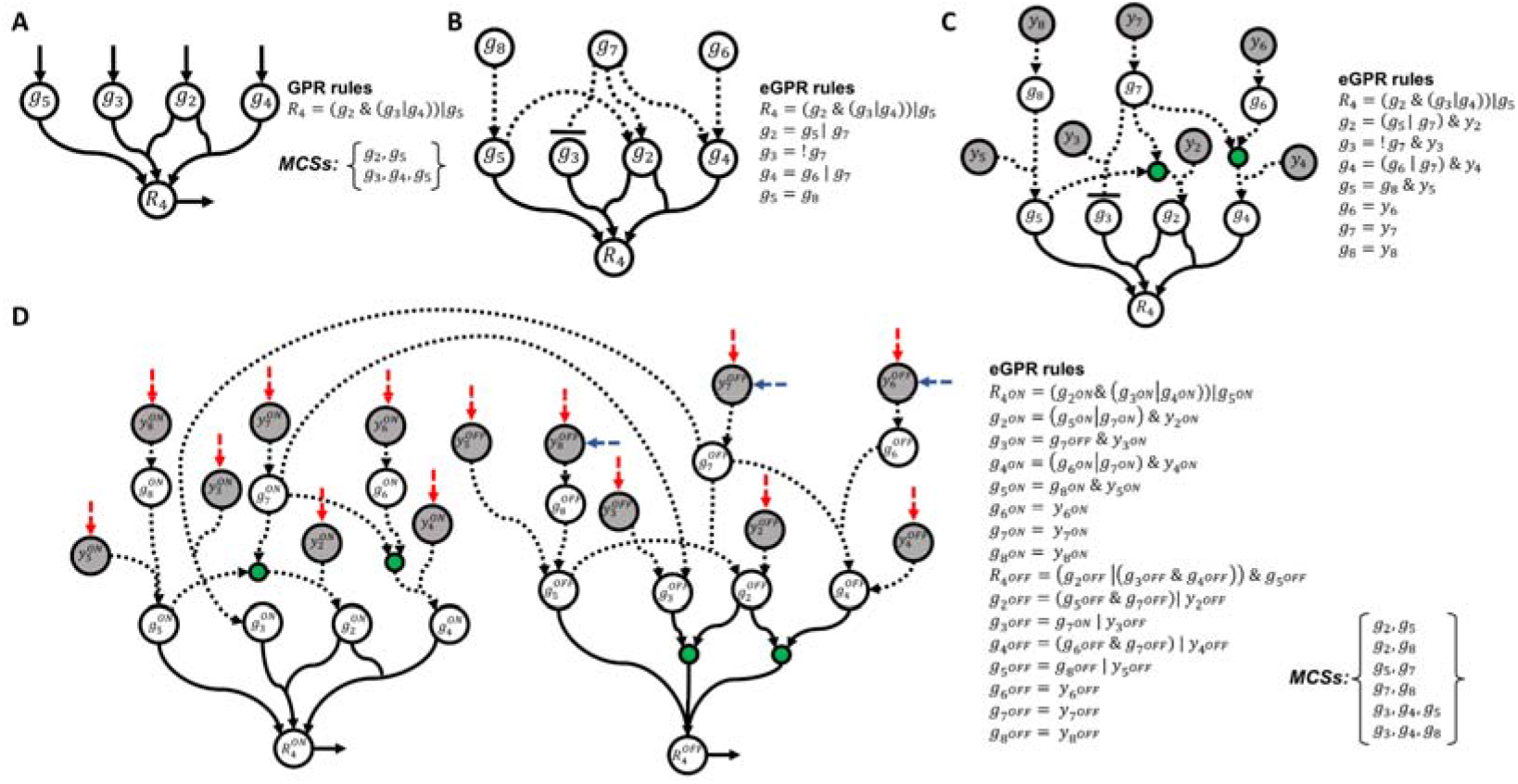
Illustration of extended GPR networks. a) Resulting GPR network for reaction 4 (*r_4_*) in Figure 1, its associated GPR rules and output MCSs; b) GPR rules including the regulation of metabolic genes involved in (*r_4_*) using Boolean equations. Three new genes are incorporated into eGPR rules: *g_6_*, *g_7_*, *g_8_*; c) Addition of auxiliary nodes *y* representing gene knockouts in Boolean equations; d) Resulting extended GPR (eGPR) network after dividing each node into two different ON/OFF nodes and including input and output exchange reactions. Regulatory interactions are represented through arcs in dashed line.

Here, we extend our computational approach to calculate gMCSs in metabolic networks that integrate linear (acyclic) regulatory pathways. In particular, we amend the *G* matrix construction algorithm to deal with the resulting acyclic Boolean networks that control metabolic reactions. The inclusion of inhibitory interactions (negation terms) in regulatory pathways requires the redefinition of our previous GPR networks and the algorithm to calculate MCSs. Figure 2b shows an example reaction that includes the regulatory information for the genes implied in its associated GPR rule. We describe below how these extended GPR (eGPR) rules are transformed into reaction networks, referred now as extended GPR (eGPR) networks, and how the MCS approach is applied to them.

### Calculation of G matrix in integrated metabolic and regulatory networks

1. *Construction of eGPR networks.* For the sake of clarity, for each target reaction *k*, denoted *R_k_*,we define *B(k)* as the subset of genes implied in its associated eGPR rules. Each of these genes, denoted *g_i_* (i=1,…,|*B(k)*|), are interconnected through their corresponding Boolean equations. We denote *L(k)* the subset of those nodes without Boolean equations (in Figure 2b, we have *g_6_*, *g_7_* and *g_8_*). Nodes in *L(k)* represent input genes for the resulting Boolean network and can freely take 0/1 values. In order to build the eGPR network for each reaction, we follow 5 different steps:

i. The Boolean equation for each gene in *B(k)* is first updated with a necessary auxiliary node *y*_i_ (*i*=1,…,| *B(k)*|), which allows us to consider the effect of gene knockouts without affecting the network upstream. The resulting Boolean network and updated eGPR rules can be found in Figure 2c. Note here that we introduce intermediate nodes (shown in green) to consider OR rules.
ii. Nodes from the Boolean network in the previous step are split into ON and OFF nodes, namely 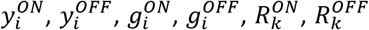 and, following the De Morgan’s laws, eGPR rules are updated. This strategy duplicates the number of nodes and interactions but negation terms disappear from the Boolean equations, which make it possible to model them as a reaction network. The resulting network is shown in Figure 2d.
iii. Addition of an input exchange reaction for nodes with no input arcs, namely 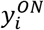 and 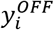. The removal of these exchange reactions represents the knockout/activation of the genes involved in our reaction network. This set of input exchange reactions is denoted *Y(k).* They are coloured red in Figure 2d.
iv. Addition of an input exchange reaction for 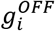 nodes such that *i* ∈ *L(k).* In general, we can reach 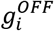 nodes from different pathways but, in the case of input genes *L(k)*, 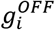 can be freely active (depending on the initial conditions). They are coloured blue in Figure 2d.
v. Addition of an output exchange reaction for 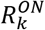 and 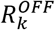, which are denoted, respectively, *r_k^ON^_* and *r_k^OFF^_* (see Figure 2d).
2. *Calculation of MCSs in eGPR networks.* eGPR networks can be modelled as a reaction system that satisfies irreversibility constraints and the mass balance equation:

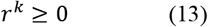

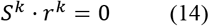

where *r^k^* denotes the flux through the artificial reactions involved in the eGPR network for the target reaction *k* and *S^k^* its associated stoichiometry matrix of dimensions *m^k^xn^k^.* In order to calculate MCSs that blocks the target reaction 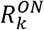, we can adapt Eq. (3) to force flux through this reaction and Eq. (4) to define the knockout space for the input exchange reactions in *Y(k)*:

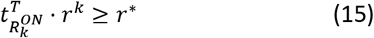

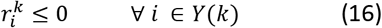

where 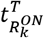 is a null vector with a 1 in the position of the target reaction 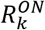. Note here that in Eq. (16) we only include input exchange reactions in *Y(k*) because they represent the decision as to whether (or not) a gene is knocked out. The knockout of 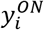 and 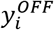 nodes are not independent, but they are coordinated in the dual problem that is presented below. The dual problem of this infeasible primal problem, Eqs. (13)–(16), takes a similar form than the one presented in Eqs. (5)–(12):

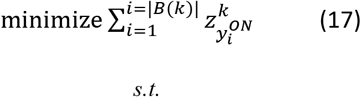

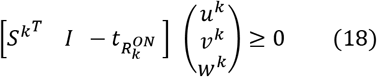

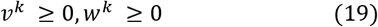

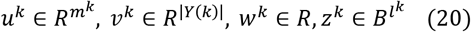

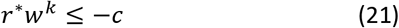

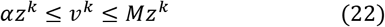

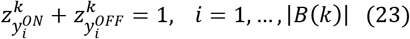

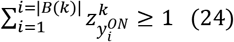

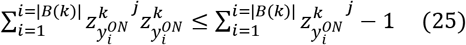 However, Eqs. (17)–(25), called MILP2, differ from MILP1 in the following points:

i. the knockout space only considers input exchange reactions associated with 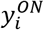 and 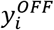, which allow us to decide which gene *i* is knocked out 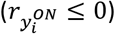 or not 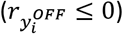 to block the target reaction;
ii. Eq. (23) forces that for each gene *i* exactly one these two constraints: 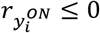 and 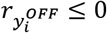 takes part in the infeasible primal problem. This constraint is specific of MILP2 and it is due to the inherent coupling between 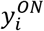 and 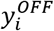 nodes. This constraint establishes that if a gene is knocked out, *i.e.* 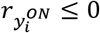, then 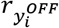 cannot be forced to be zero, and vice versa.
iii. the objective function, Eq. (17), minimizes the number of knockouts of input exchange reactions associated with 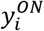, since they represent gene knockouts 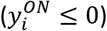. The same logic applies to the solution elimination constraint in Eq. (25).
iv. We force the optimal solution to involve at least one gene knockout in Eq. (24). MILP2, Eqs. (17)–(25), allows us to enumerate MCSs for eGPR networks. Figure 2d shows the resulting MCSs for the eGPR network of reaction 4 in Figure 1.
3. *Calculation of G matrix.* Using as input data the GPR rules and regulation available for a specific metabolic network, MILP2 is applied to each different reaction. For illustration, Figure 3a shows the example metabolic network in Figure 1a, but additionally including the regulation for some of the metabolic genes involved (eGPR rules). Figure 3b shows the resulting MCSs for each target reaction after applying MILP2 to its associated eGPR network. MCSs for different reactions are then integrated in order to build *G* and *F* matrix (see Figure 3b). We have developed a MATLAB function for building the *G* matrix in integrated metabolic and regulatory models, called *‘buildGmatrix_iMRmodel’*, which can be integrated in the COBRA toolbox **(*Heirendt* et al., 2019)**. Note here that as the size of *G* matrix increases with the addition regulatory interactions, we have conducted several improvements in this function, reducing up to 3 times the computation time with respect our previous implementation.

**Figure 3.**
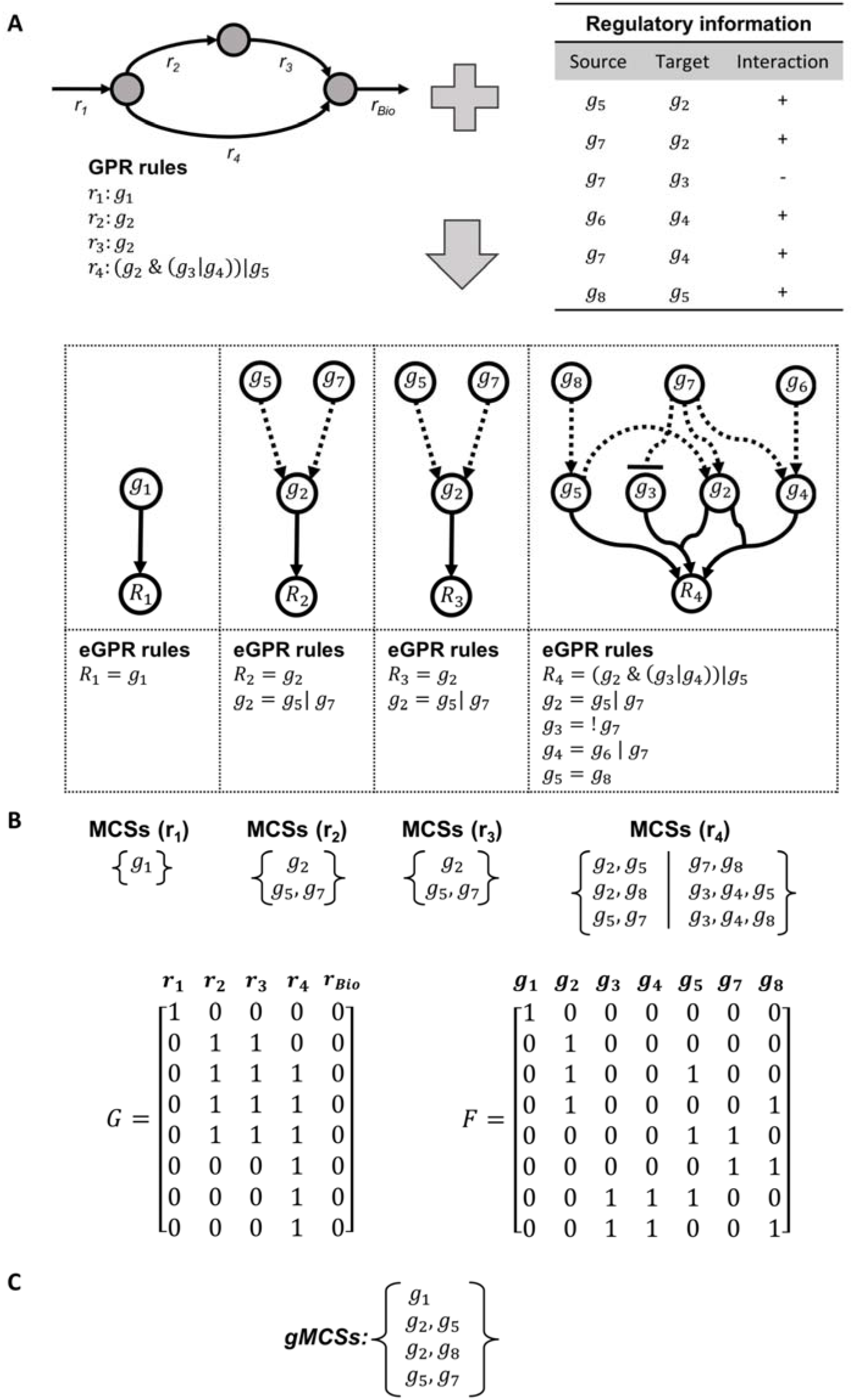
Illustration of gMCSs in integrated metabolic and regulatory models. A) Example integrated metabolic and regulatory model that extends the metabolic network in Figure 1. B) Resulting MCSs for each target reaction after applying MILP2 to its associated eGPR network. In addition, *G* and *F* matrices are provided. C) Resulting gMCSs for this toy example integrated network.

Once the *G* matrix has been obtained, the list of gMCSs can be calculated using the function *‘calculateGeneMCS’*, also presented in Apaolaza et al., 2019, which makes use of MILP1. The resulting gMCSs for our toy example can be found in Figure 3c.

### Definition of regulation layers in metabolic models

In order to define the regulation layer of the metabolic network under study, we first find, using different databases (see Results section), signed interactions for each metabolic gene involved in GPR rules. Then, we create a new Boolean equation that integrates the identified interactions for each metabolic gene using ‘OR’ operators, as observed in Figure 3A, leading to eGPR rules.

As noted above, the methodology developed in this work (MILP2) is not able to deal with cyclic behaviors that are common in Boolean networks. For that reason, at the time of adding a regulatory layer, we must check that there are no cycles in the resulting eGPR network. This is done by solving the following linear programming problem (LP1) for each reaction *R_k_*:

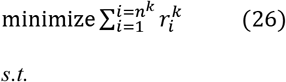

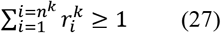

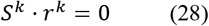

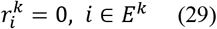

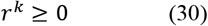

where *E^k^* is the subset of input and output exchanges in the eGPR network for reaction *R_k_.*

If we delete input and output exchanges fluxes with Eq. (29), LP1 is only feasible in the case we have cycles in the eGPR network, otherwise the solution is infeasible. Once it is tested that the eGPR network does not present cycles (LP1 is infeasible), the regulatory layer is added to the model. Note here that adding a layer involves including more genes to the model which can be regulated by other genes. Therefore, we can search for all the regulatory interactions related to the genes added in the previous layer and insert new genes to the network as explained above. Then, the absence of cycles is checked and the layer is added. This process can be repeated as many times as layers are desired to be added to the model. Supplementary Figure 1 shows the toy example in Figure 1 with one, two and three regulation layers.

### Implementation

For the different studies conducted in the Results section, we used the University of Navarra’s computing cluster, limiting to 8 cores and 8 GB of RAM (in 2 cases we required 32 GB of RAM). A time limit of 5 minutes was set for each solution derived from the function *‘CalculateGeneMCS’.* MATLAB and The COBRA toolbox was used to implement the function ‘buildGmatrix_iMRmodel’, with help of IBM Ilog Cplex for the underlying MILP model.

## Results

In order to assess the performance of our novel approach, we integrated the extensively curated and most recent published metabolic network of human cells, Human1 (Robinson *et al.*, 2020) (v1.10.0), with the protein-protein interaction network of Omnipath (Türei *et al.*, 2016) (accessed online 2021-09-29) (OmnipathR, v.3.0.4) (Valdeolivas A, Turei D, 2019), the gene regulatory network of signed transcription factors Dorothea (Garcia-Alonso *et al.*, 2019) and the manually curated database of human transcriptional regulatory networks TRRUST (Han *et al.*,2018). In summary, we built 3 different integrated models: Human1 + Omnipath (Human1-O); Human1 + Dorothea (Human1-D), Human1 + TRRUST (Human1-T). We present below the analysis of identified gMCSs for these integrated models with single and multiple regulatory layers.

### Analysis of gMCSs in single-layer integrated metabolic and regulatory models

Table1 shows the details for the different models mentioned above including a unique layer of regulatory interactions for each metabolic gene (see Methods section). The addition of the regulatory layer significantly increases the number of genes in the 3 cases, being Human1-O1 the one with the highest increase. However, we obtained the most complex *G* matrix and highest computation time with Human1-D1. It can be observed that the computation time scales linearly with the number of rows of *G* matrix (Pearson’s correlation=0.99, p-value=0.007672).

**Table 1.** Summary of single-layer integrated metabolic and regulatory models and computed gMCSs. Computation time is given in seconds (s). Abbreviations: *‘Human1-O1’*: integrated model with Human1 and Omnipath with one regulatory layer; *‘Human1-D1’*: integrated model with Human1 and Dorothea with one regulatory layer; *‘Human1-T1’:* integrated model with Human1 and TRRUST with one regulatory layer. In the column *‘Number of gMCSs (length≤5)’*, the number in parenthesis is the number of gMCSs arising from the addition of the regulatory layer.

| Model | Number of genes | G matrix dimension | G matrix computation time (s) | Number of gMCSs (length ≤ 5) |
| --- | --- | --- | --- | --- |
| Human1 | 2,975 | 1,706x10,342 | 314 | 9881 |
| Human1-O1 | 3,419 | 2,407x10,342 | 634 | 12751 (2870) |
| Human1-D1 | 3,072 | 5,233x10,342 | 1,332 | 11020 (1139) |
| Human1-T1 | 3,214 | 2,724x10,342 | 634 | 10557 (676) |

For each model, we calculated gMCSs until length 5 that block biomass production (Table 1). 9881 gMCSs were identified for Human1. All of them were included in our 3 integrated models (Supplementary Figure 2); however, we found 4686 new gMCSs: 2870 in Human1-O1, 1177 in Human1-D1 and 677 in Human1-T1. We observed that the new subset of gMCSs identified strongly depends on the regulatory database employed and shows limited overlap (Supplementary Figure 2).

Given the differences found in the different integrated models, we compared their capacity to predict essential genes in cancer, following the gMCS approach recently developed in our group (Valcárcel *et al.*, 2022). We used as a gold-standard the genome-wide CRISPRi experiments from 5 cancer cell lines published by Hart and colleagues (Hart *et al.*, 2015), referred them to as Hart2015. In Valcárcel et al., 2022, a gene is classified as essential in a sample if it is the unique highly expressed gene in at least one gMCS and the rest of genes of that gMCS are lowly expressed. For the definition of highly and lowly expressed genes for each sample, we applied the *gmcsTH5* threshold presented in that work (Valcárcel *et al.*, 2022) and gene expression data from CCLE (Ghandi *et al.*, 2019). For consistency, *gmcsTH5* was derived for each sample using the gMCSs calculated for Human1 and applied to the rest of the models. Once the list of essential genes per cell line and per integrated model were computed, we compared them with the essentiality scores of Hart2015. We determined the number of true positives (TPs) and false positives (FPs), as well as the positive predictive value (PPV), which is the ratio TPs to all of the genes that were defined as positive (TP + FP) (Figure 4).

**Figure 4.**
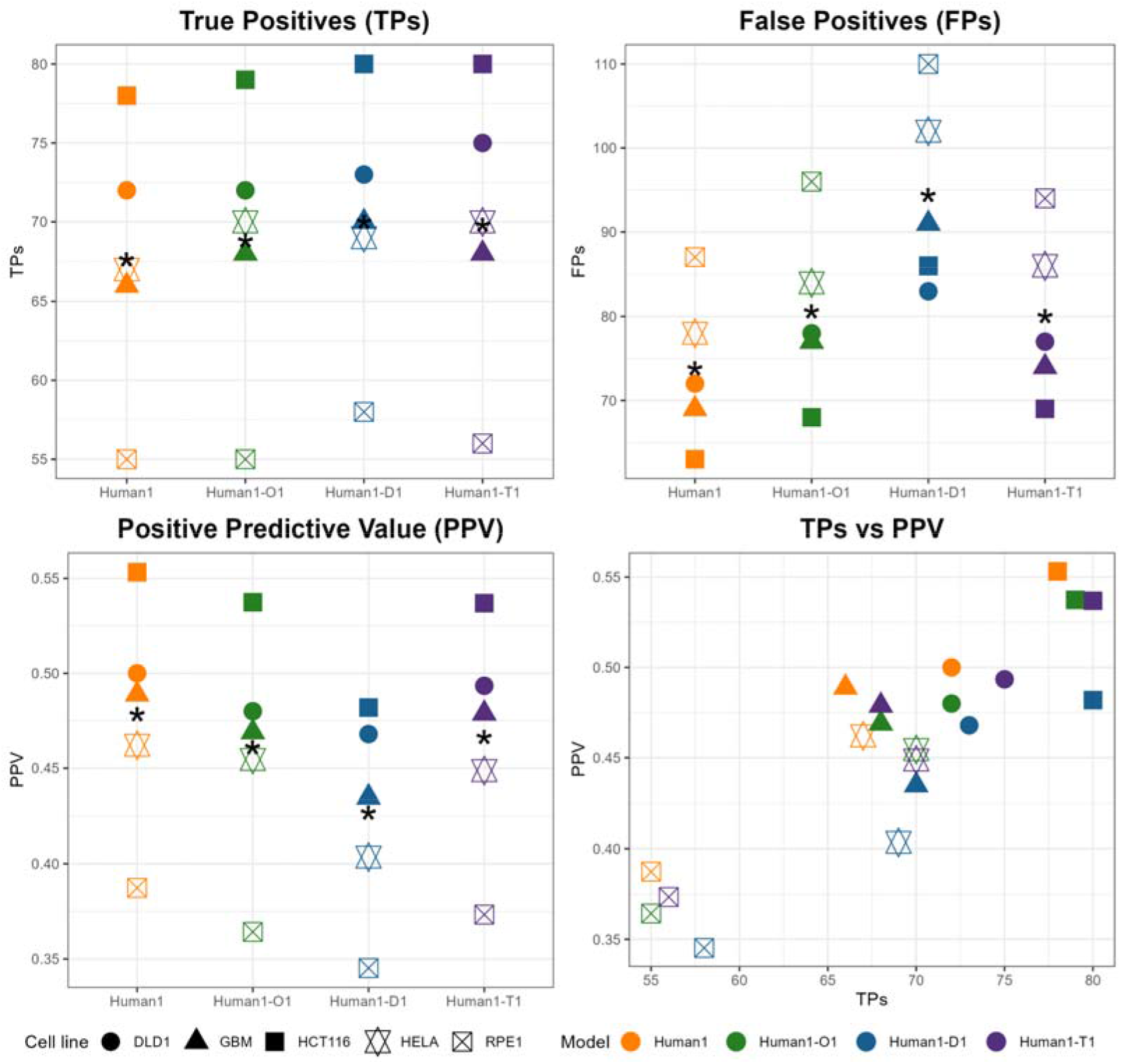
Gene essentiality comparison between Human1 and single-layer integrated metabolic and regulatory models. True Positives (TPs), False Positives (FPs), Positive Predictive Value (PPV) and TPs vs PPV arising from our different models (Human1, Human1-O1, Human-D1, Human1-T1) using the essentiality data presented in Hart2015 as a gold-standard. Asterisk * represents the mean value for the 5 cell lines considered.

As shown in Figure 4A, the addition of a regulatory layer involves a significant increase in the number of TPs. However, FPs also rise under the addition of the regulatory layer and, for this reason, the PPV of the integrated models is slightly lower than in Human1. Interestingly, as it is shown in the plot TPs vs PPV, although the PPV value decreases by the addition of the regulatory layer, the integrated models always dominate Human1 in terms of TPs. Dorothea seems to be the regulatory database that leads to the detection of more TPs, but it is also the one with the highest value of FPs and, so, the lowest PPV of all the models. TRRUST database seems to present the best proportion of TPs and FPs in relation to Human1. A similar conclusion was obtained for DepMap data (Ghandi *et al.*, 2019; Meyers *et al.*, 2017) (Supplementary Figure 3).

Each regulatory database led to the detection of specific subsets of essential genes. For example, in the cell line HELA, we found the same 145 metabolic essential genes in all the models; 26 new essential genes with Human1-D1, among which 4 are shared with Human1-T1 and 1 with Human1-O1; 9 new essential genes with Human1-O1, among which 1 is shared with Human1-D1; and 11 new essential genes in Human1-T1, among which 4 are shared with Human1-D1 (Supplementary Figure 4). In addition, new essential genes are transcription factors but also metabolic enzymes. An example in this cell line is TXN2. In Human1, it appears in a unique gMCSs of length 2: TXN & TXN2. TXN is expressed in HELA, and, for that reason, TXN2 is not predicted as essential in Human1. However, in Human1-T1, TXN2 appears in 2 gMCSs: {TXN2 & TXN}, {TXN2 & PPARD}. In HELA, the gene PPARD is not expressed and, thus, TXN2 is predicted as an essential gene.

### Analysis of gMCSs in multiple-layer integrated metabolic and regulatory models

Table2 shows the details for the different integrated models including a 1, 2 and 3 layers of regulatory interactions for each metabolic gene (see Methods section). The addition of multiple layers has particularly an impact in Human1-O, which involves 4371 genes in the third layer (Human1-O3). In Human1-D and Human1-T, the impact of multiple layers is moderate and the third layer seems irrelevant, because they are smaller databases. In addition, we obtained the most complex *G* matrix and highest computation time with Human1-O2 and Human1-O3 (see Table 2). As it was found in the single layer analysis, the computation time scales linearly with the number of rows of *G* matrix (Pearson’s correlation=0.99, p-value=4.646e-08).

**Table 2.** Summary of multiple-layer integrated metabolic and regulatory models and computed gMCSs. Results correspond to models including 1, 2 and 3 regulatory layers. Computation time is given in seconds (s). In the column *‘Number of gMCSs (length≤3)’*, the number in parenthesis is the number of gMCSs arising from the addition of the regulatory layer. Abbreviations: *‘Human1-O1’*: integrated model with Human1 and Omnipath with one regulatory layer; *‘Human1-O2’*: integrated model with Human1 and Omnipath with two regulatory layers; *‘Human1-O3’*: integrated model with Human1 and Omnipath with three regulatory layers; *‘Human1-D1’*: integrated model with Human1 and Dorothea with one regulatory layer; *‘Human1-D2’*: integrated model with Human1 and Dorothea with two regulatory layers; *‘Human1-D3’*: integrated model with Human1 and Dorothea with three regulatory layers; *‘Human1-T1’*: integrated model with Human1 plus TRRUST and one regulatory layer; *‘Human1-T2’*: integrated model with Human1 plus TRRUST and two regulatory layers; *‘Human1-T3’*: integrated model with Human1 and TRRUST with three regulatory layers.

| Model | Number of genes | G matrix dimension | G matrix computation time (s) | Number of gMCSs ($length \leq 3$) |
| --- | --- | --- | --- | --- |
| Human1 | 2,975 | 1,706x10,342 | 314 | 249 |
| Human1-O1 | 3,419 | 2,407x10,342 | 634 | 269 (20) |
| Human1-O2 | 4,371 | 19,791x10,342 | 8,623 | 621 (372) |
| Human1-O3 | 4,525 | 20,494x10,342 | 9,308 | 441 (192) |
| Human1-D1 | 3,072 | 5,233x10,342 | 1,332 | 292 (43) |
| Human1-D2 | 3,081 | 6,077x10,342 | 2,078 | 296 (47) |
| Human1-D3 | 3,081 | 6,178x10,342 | 2,154 | 298 (49) |
| Human1-T1 | 3,214 | 2,724x10,342 | 634 | 260 (11) |
| Human1-T2 | 3,358 | 6,534x10,342 | 1,581 | 266 (17) |
| Human1-T3 | 3,382 | 8,438x10,342 | 2,168 | 291 (42) |

For each model, we calculated gMCSs until length 3 that block biomass production (Table 2). 248 gMCSs were identified for Human1. All of them were included in our 9 integrated models (Supplementary Figure 5). Human1-O2 and Human1-O3 increased significantly the number of gMCSs. This is not observed either in Human1-D2 and Human1-D3 or in Human1-T2 and Human1-T3, in line with the dimension of *G* matrix. Again, we observed that the new subset of gMCSs identified strongly depends on the regulatory database employed and the intersection between databases is limited (Supplementary Figure 5). Note here that the memory requirement in Human1-O2 and Human-O3 exceeded 16Gb, which made it impossible performing the computation of gMCSs in a standard computer.

We conducted the same gene essentiality analysis shown above for multiple-layer integrated models (Figure 5). In the case of Human1-O, the number of TPs increased to significantly lower rate than the number of FPs after adding the second and third layer (Human1-O2 and Human1-O3), which substantially decreased PPV in comparison with Human1. In the case of Human1-D and Human1-T, the effect of multiple layers seems less relevant. We obtained similar conclusions with DepMap data (Supplementary Figure 6).

**Figure 5.**
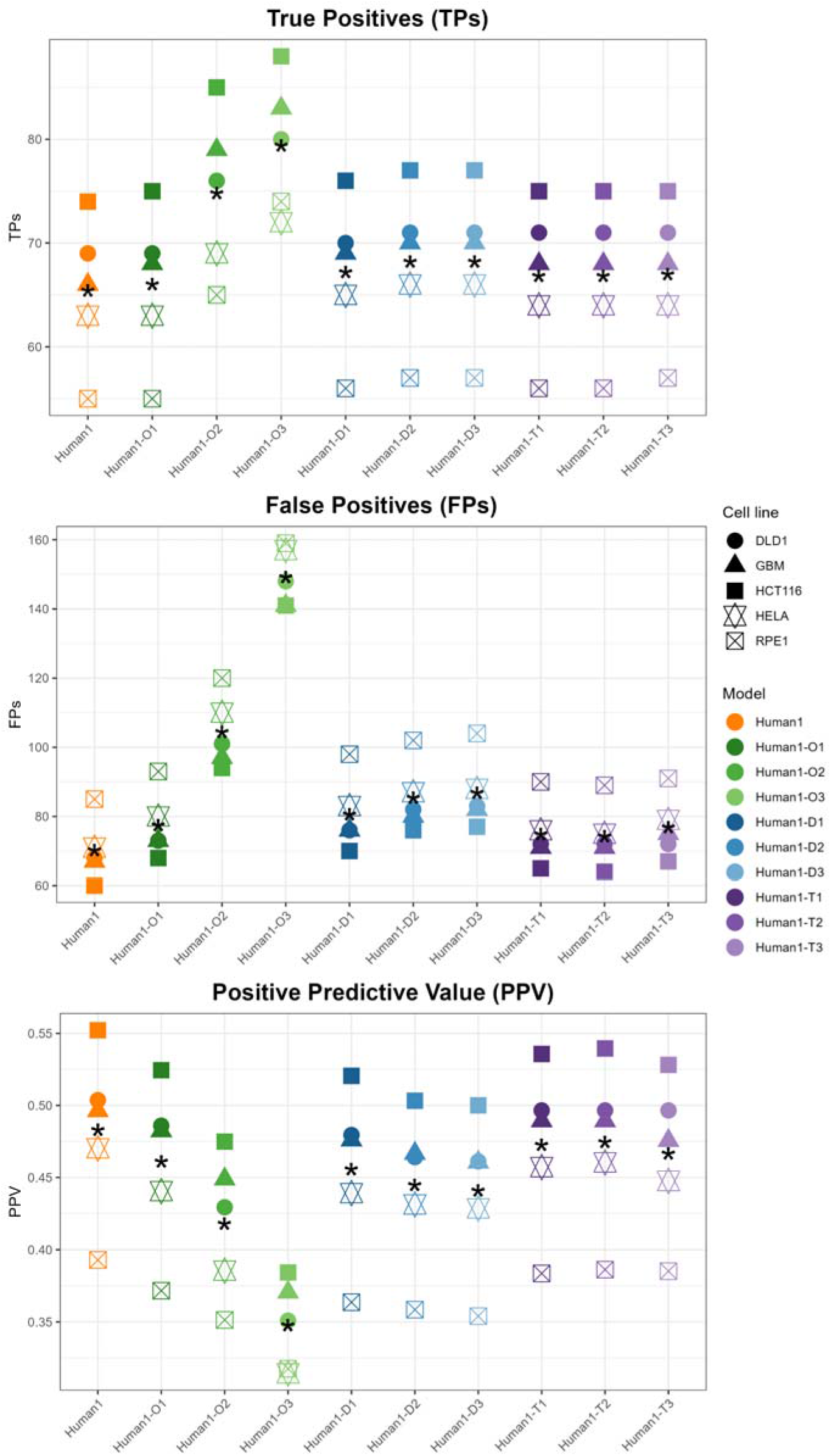
Gene essentiality comparison between Human1 and multiple-layer integrated metabolic and regulatory models. True Positives (TPs), False Positives (FPs), Positive Predictive Value (PPV) and TPs vs PPV arising from our different models (Human1, Human1-O1, Human1-O2, Human1-O3, Human-D1, Human-D2, Human-D3, Human1-T1, Human1-T2, Human1-T3) using the essentiality data presented in Hart2015 as a gold-standard. Asterisk * represents the mean value for the 5 cell lines considered.

## Discussion

The integration of genome-scale metabolic and regulatory models has received considerable attention in the literature. Most algorithms aim to integrate regulatory networks to refine the prediction of metabolic fluxes (Chandrasekaran and Price, 2010; Marmiesse et al., 2015; Wang et al., 2017). However, the identification of synthetic lethals from these integrated models have been little explored. Early approaches rely on pathway enumeration, which are not tractable for genome-scale models (Jungreuthmayer and Zanghellini, 2012). Here, using the concept of gMCSs, we present the first approach to address this issue in large-scale networks.

The search of synthetic lethals in these integrated metabolic and regulatory models poses different challenges. Complex regulatory networks, represented here by Boolean networks, involve negation terms and cycles, which are not present in metabolic GPR rules. In this work, we partially address this problem and adapt our previous gMCS formulation to integrate linear regulatory pathways. The consideration of regulatory cycles in our approach is pendant and it will be addressed in future works.

Our novel gMCS approach was applied to predict synthetic lethality in human cells. To that end, we integrated the most recent draft metabolic model of human cells, Human1 (Robinson *et al.*,2020), with Omnipath (Türei *et al.*, 2016), Dorothea (Garcia-Alonso *et al.*, 2019) and TRRUST (Han *et al.*, 2018). For each regulatory network, we built a different integrated model and effectively enumerated gMCSs. In particular, we present results for these integrated models under a single (gMCSs up to length 5) and multiple (gMCSs up to length 3) regulatory layers. Our gMCS approach was effective in all the cases considered, including networks involving more than 4500 genes, which opens the door to incorporate other regulatory layers.

We compared the performance of the different integrated models with gene essentiality data from human cancer cell lines. As shown in Figure 5, we found that models with single regulation layers seem more accurate than models with multiple regulation layers. This is particularly true in the case of Omnipath, where models with multiple regulation layers substantially increased the number of genes and resulting gMCSs, but the accuracy in the essentiality predictions is significantly lower. This lack of accuracy may be caused by different reasons: 1) Omnipath integrates different sources of information with a different quality of annotation; 2) the bias introduced in our predictions for not considering cyclic interactions; 3) annotation errors are propagated along multiple regulatory layers. In the case of Dorothea and TRRUST, considerably smaller databases than Omnipath, the negative effect of multiple regulatory layers is less relevant and the differences between single and multiple regulatory layers are small in terms of number of genes, gMCSs and essential genes.

We also compared the gene essentiality predictions obtained from the different regulatory networks. A significant number of new essential genes was obtained for the different integrated models. However, we found that TRRUST returned the most accurate results. Interestingly, the overlap between TRRUST and the other 2 databases is low, which suggests that the integration of them could lead to more accurate regulatory models. In fact, we assessed an integrated model with TRRUST and Dorothea (deleting contradicting interactions), finding that gene essentiality predictions are slightly better than exclusively TRRUST (Supplementary Figure 7). Thus, integrating and comparing different regulatory databases in the light of gene essentiality predictions could be an interesting future direction.

We analysed in detail essential genes and synthetic lethals obtained with TRRUST. We found 5 new essential genes for all cell lines (gMCSs of length 1): E2F1, KLF5, NR1H4, SP1 and SREBF2. We found extensive literature supporting our predictions for E2F1 and KLF5 (Wu et al., 2001, Dong and Chen, 2009). In addition, we found that SP1 is over-expressed in most tumours and an attractive target for cancer cells (Vizcaíno *et al.*, 2015), and that SREBF2 is essential for tumour growth and initiation in colon cancer (Wen *et al.*, 2018).

Regarding to the new synthetic lethals and context-specific essential genes obtained with TRRUST, a summary list can be found in Supplementary Table 1. Interestingly, we predicted two essential metabolic genes that were not captured by Human1: PISD and TXN2, which shows the potential of our integrated approach to complement previous predictions. In particular, PISD was predicted essential in HCT116 and HELA cell lines, in line with Bellance and collegues (Bellance *et al.*, 2020), where they demonstrated that doxorubicin inhibits PISD and induces cell death in HELA cells. Similarly, TXN2 was predicted essential in DLD1 and HELA cells, in agreement with the work presented in Zhang et al., 2011, where they proved that knockdown of TXN2 caused a significant decrease of cell viability in HELA. On the other hand, we predicted the essentiality of CREB1 in all cell lines in Hart2015. CREB1 is a transcription factor that comprises a synthetic lethal with ACACB, a metabolic gene implied in fatty acid biosynthetis and biotin metabolism (Supplementary Figure 8). ACACB is lowly expressed in all the cell lines, and so, the inhibition of CREB1 leads to cell death. The literature is also supporting of our prediction, since Fang et al., 2016 showed that the downregulation of CREB1 is lethal in HCT116. This synthetic lethal shows again the functional interaction between the metabolic and regulatory layers.

Overall, our novel gMCS approach opens avenues to predict mechanistically synthetic lethal interactions between metabolic and regulatory genes. The computational and functional (biological) analysis presented here shows that our tool can be robustly used to study the regulation of cancer metabolism and associated dependencies.

## Data availability

The authors confirm that the data supporting the findings of this study are available within the article and its supplementary material.

## Code availability

MATLAB code is available in https://github.com/nbarrenac/cobratoolbox/tree/extended_gMCS_2022/src/analysis/gMCS.

## Acnowledgements

This work was supported by the Minister of Economy and Competitiveness of Spain [PID2019-110344RB-I00, F.J.P.], PIBA Programme of the Basque Government [PIBA_2020_01_0055, F.J.P.], ERANET program ERAPerMed [MEET-AML, F.P.], Elkartek programme of the Basque Government [KK-2022/00045, F.J.P.] and Fundación Ramon Areces [PREMMAN, F.J.P.]. N.B. received his salary from Basque Government predoctoral grant [PRE_2021_2_0025]. The funders had no role in study design, data collection and analysis, decision to publish, or preparation of the manuscript

## Author contributions

F.J.P. conceived this study. N.B., L.V.V., I.A. and F.J.P. developed the mathematical model to calculate gMCSs in integrated metabolic and regulatory network models. N.B., L.V.V. and D.O. carried out the computational implementation in MATLAB. N.B., L.V.V., I.A. and F.J.P. designed and performed gene essentiality analysis of cancer cell lines. All authors wrote, read, and approved the manuscript.

