## Supplementary Material for "Synthetic lethality in large-scale integrated metabolic and regulatory network models of human cells"

**A**

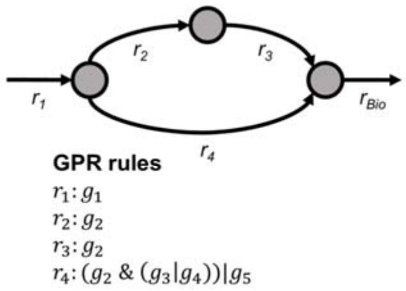


**Regulatory information**

| Source | Target | Interaction |
|---|---|---|
| $g_5$ | $g_2$ | + |
| $g_7$ | $g_2$ | + |
| $g_7$ | $g_3$ | - |
| $g_6$ | $g_4$ | + |
| $g_7$ | $g_4$ | + |
| $g_8$ | $g_5$ | + |
| $g_9$ | $g_7$ | - |
| $g_4$ | $g_8$ | + |
| $g_{10}$ | $g_8$ | + |
| $g_6$ | $g_{10}$ | + |
| $g_9$ | $g_{10}$ | + |

**B**

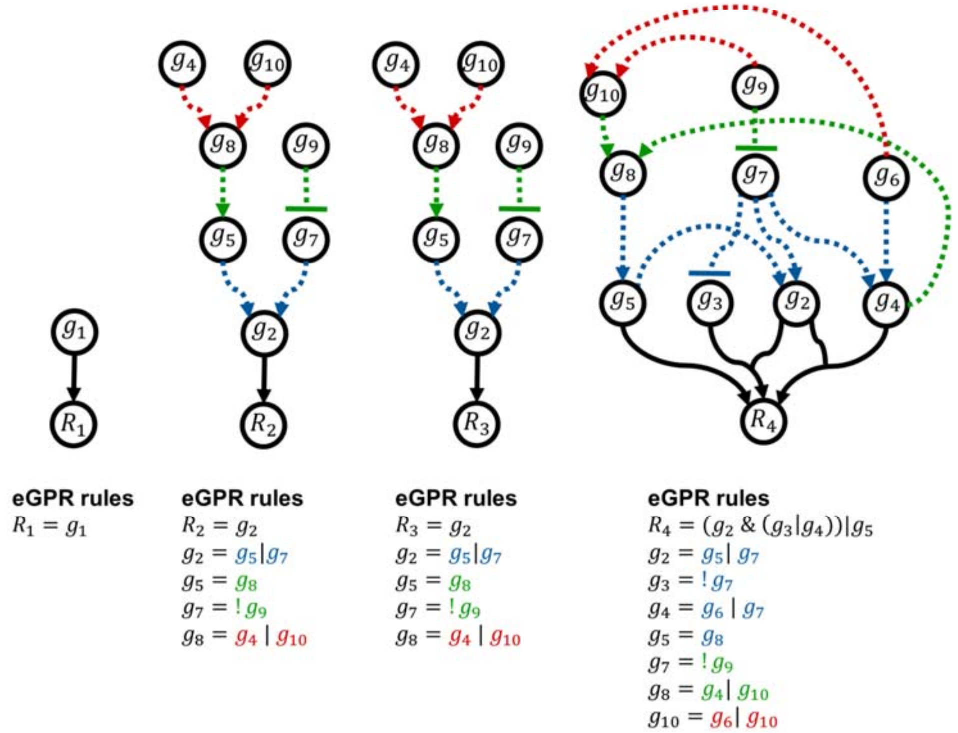


**Supplementary Figure 1. Example network in Figure 1 with 1, 2 and 3 regulatory layers.** a) Example network in Figure 1 and associated regulatory information. b) Systematic addition of regulatory information. Arrows in blue represent the first regulatory layer; arrows in green represent the second regulatory layer and arrows in red represent the third regulatory layer.

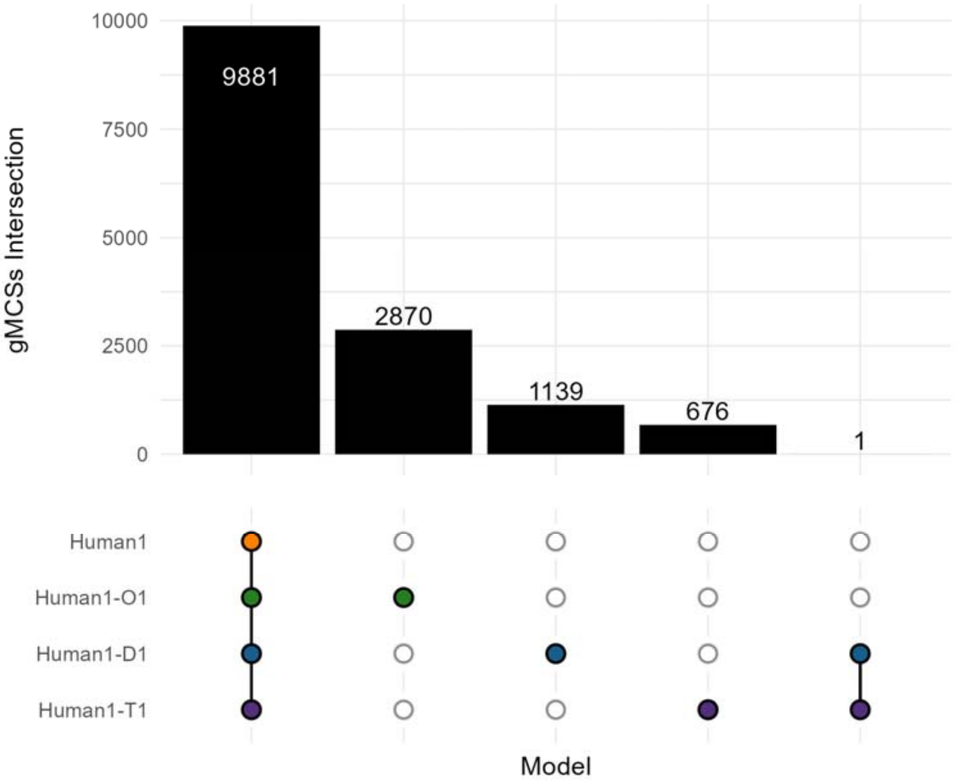


**Supplementary Figure 2. Analysis of gMCSs obtained from single-layer integrated metabolic and regulatory models.** Upsetplot representing the intersections of gMCSs until length 5 calculated from different integrated models: Human1, Human1-O1, Human-D1, Human1-T1. Abbreviations: '*Human1-O1'*: integrated model with Human1 and Omnipath with one regulatory layer; '*Human1-D1*': integrated model with Human1 and Dorothea with one regulatory layer; '*Human1-T1*': integrated model with Human1 and TRRUST with one regulatory layer.

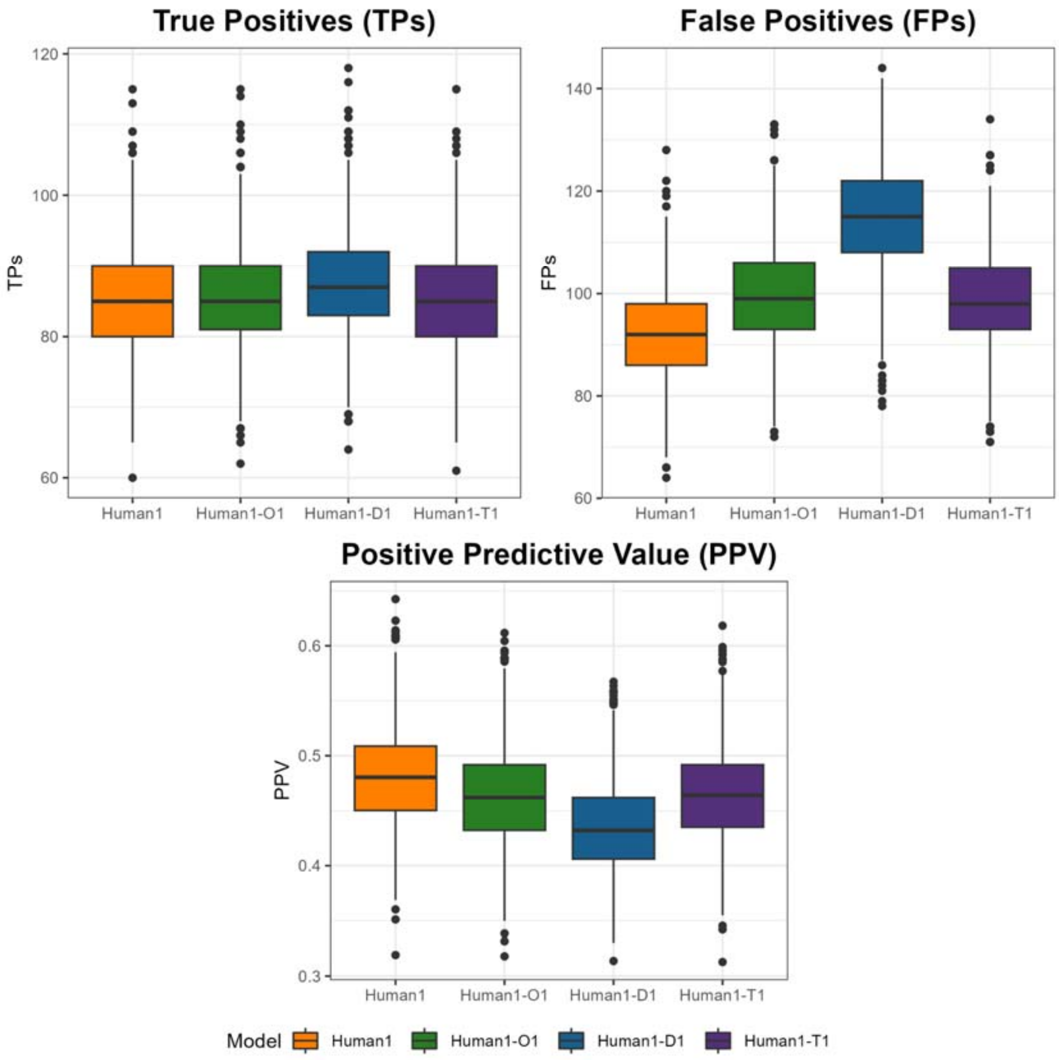


**Supplementary Figure 3. Gene essentiality comparison between Human1 and single-layer integrated metabolic and regulatory models using data from DepMap.** True Positives (TPs), False Positives (FPs) and Positive Predictive Value (PPV) arising from our different models (Human1, Human1-O1, Human-D1, Human1-T1) using the essentiality data presented in DepMap.

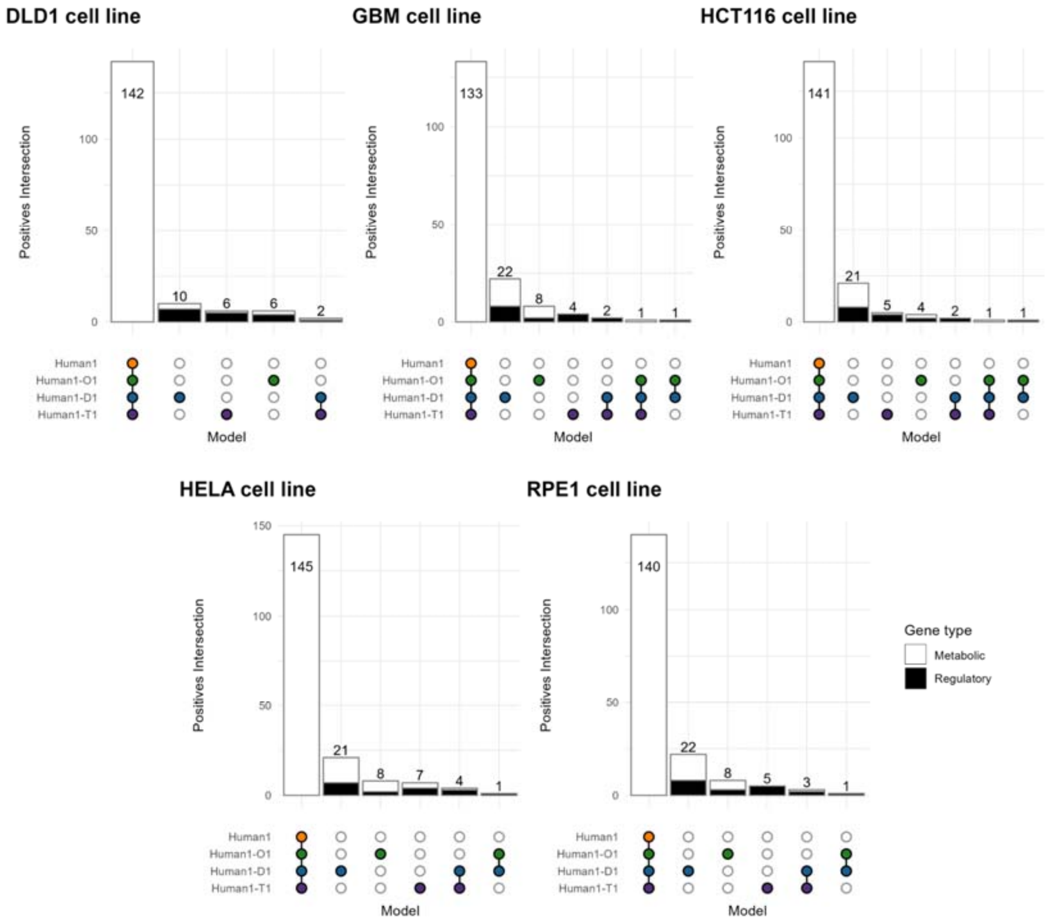


**Supplementary Figure 4. Predicted number of essential genes in each cell line of Hart2015 with Human1 and single-layer integrated metabolic and regulatory models.** Upsetplot representing the intersection of the predicted essential genes in Hart2015 cell lines (DLD1, GBM, HCT116, HELA, RPE1) derived from the gMCSs obtained from the different models analyzed: Human1, Human1-O1, Human-D1, Human1-T1.

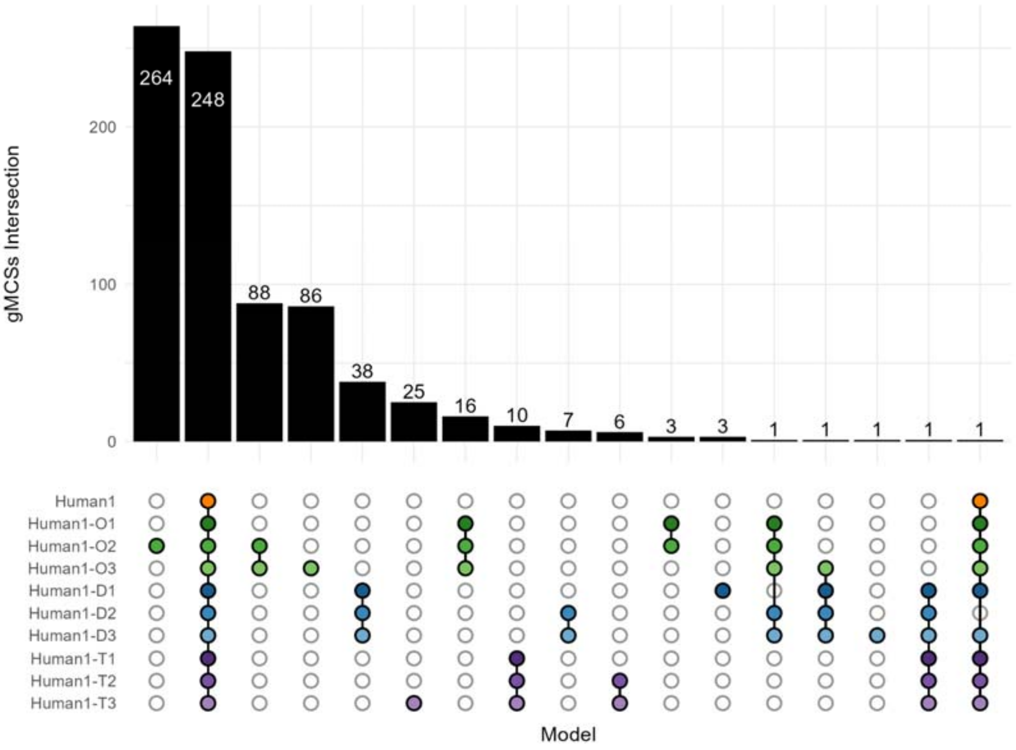


**Supplementary Figure 5. Analysis of gMCSs obtained from multiple-layer integrated metabolic and regulatory models**. Upsetplot representing the intersection of the gMCSs until length 3 calculated in our different models: Human1, Human1-O1, Human1-O2, Human1-O3, Human-D1, Human-D2, Human-D3, Human1-T1, Human1-T2, Human1-T3. Abbreviations: 'Human1-O1': integrated model with Human1 and Omnipath with one regulatory layer; 'Human1-O2': integrated model with Human1 and Omnipath with two regulatory layers; 'Human1-O3': integrated model with Human1 and Omnipath with three regulatory layers; 'Human1-D1': integrated model with Human1 and Dorothea with one regulatory layer; 'Human1-D2': integrated model with Human1 and Dorothea with two regulatory layers; 'Human1-D3': integrated model with Human1 and Dorothea with three regulatory layers; 'Human1-T1': integrated model with Human1 plus TRRUST and one regulatory layer; 'Human1-T2': integrated model with Human1 plus TRRUST and two regulatory layers; 'Human1-T3': integrated model with Human1 and TRRUST with three regulatory layers.

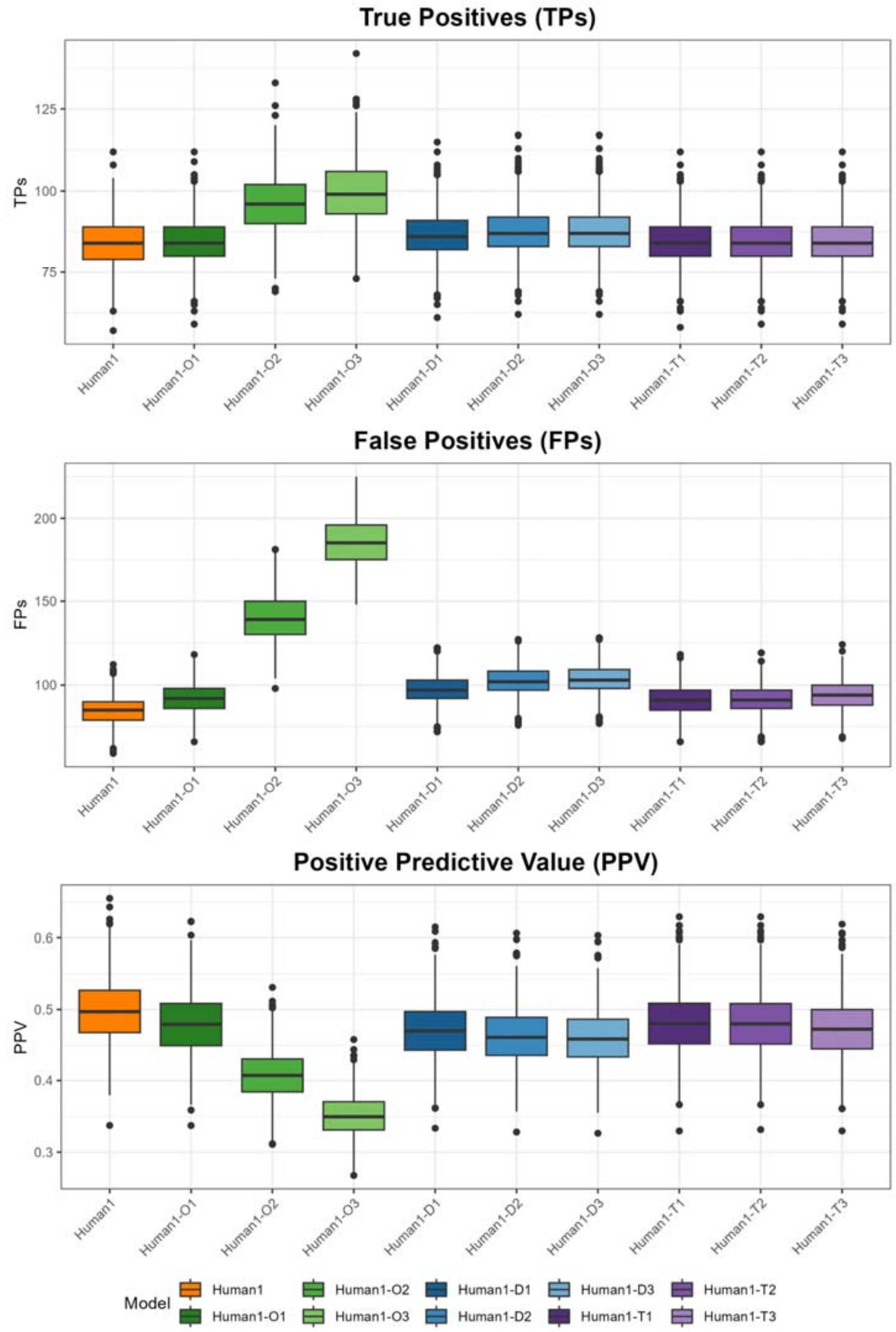


**Supplementary Figure 6. Gene essentiality comparison between Human1 and multiple-layer integrated metabolic and regulatory models using data from DepMap.** True Positives (TPs), False Positives (FPs) and Positive Predictive Value (PPV) arising from our different models (Human1, Human1-O1, Human1-O2, Human1-O3, Human-D1, Human-D2, Human-D3, Human1-T1, Human1-T2, Human1-T3) using the essentiality data presented in DepMap.

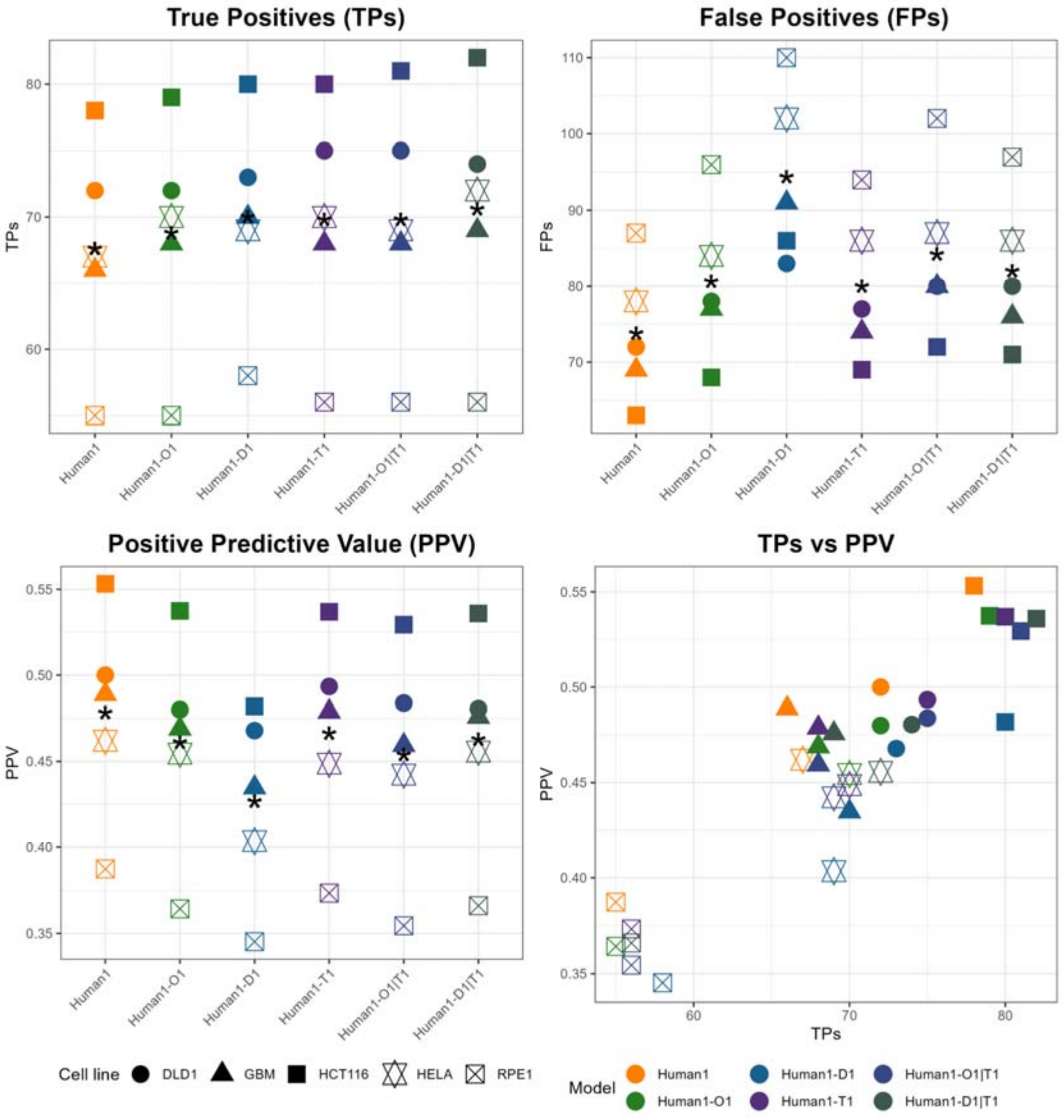


**Supplementary Figure 7. Effect of the combination of different regulatory databases in the gene essentiality comparison with Hart2015.** True Positives (TPs), False Positives (FPs) and Positive Predictive Value (PPV) arisings from our different single-layer integrated metabolic and regulatory models considered: Human1, Human-D1, Human1-T1, Human1-O1|T1, which combines Omnipath and TRRUST for the regulation layer, and Human1-D1|T1, which combines Dorothea and TRRUST for the regulation layer. Asterisk * represents the mean value for the 5 cell lines considered.

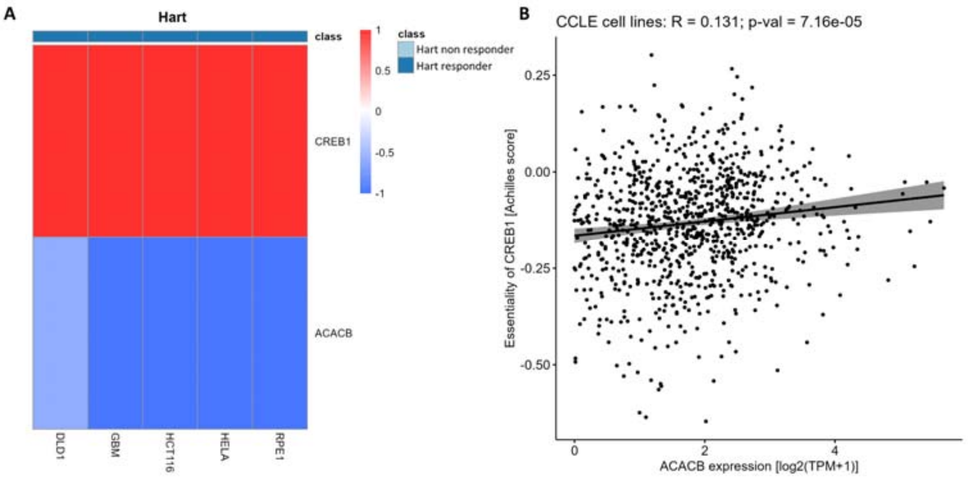


**Supplementary Figure 8. Prediction of essentiality of CREB1 in Hart2015 and correlation analysis in DepMap cell lines.** A) Expression of the genes CREB1 and ACACB, which comprise a new gMCS that is predicted in Human1-T1. CREB1 is predicted essential in all the cell lines of Hart2015 because ACACB is not expressed. B) Correlation between the essentiality of CREB1 (CRISPR knockout screen data from DepMap) and the expression of ACACB in log2(TPM+1).

**Supplementary Tables**

**Supplementary Table 1.** Summary list of the new synthetic lethals and context-specific essential genes obtained with Human1-T1 (single-layer integrated model with Human1 and TRRUST).

| Gene | Gene type | gMCS | Cell line | Literature |
|---|---|---|---|---|
| E2F1 | Regulatory | {E2F1} | ALL | (Wu *et al.*, 2001) |
| KLF5 | Regulatory | {KLF5} | ALL | (Dong and Chen, 2009)<br>(Takagi *et al.*, 2020) |
| NR1H4 | Regulatory | {NR1H4} | ALL | |
| SP1 | Regulatory | {SP1} | ALL | (Vizcaíno *et al.*, 2015)<br>(Zhao *et al.*, 2013)<br>(Lee *et al.*, 2015) |
| SREBF2 | Regulatory | {SREBF2} | ALL | (Wen *et al.*, 2018) |
| ACAT2 | Metabolic | {ACAT2, EHHADH, TMEM91, HPD, HMGCS2} | DLD1, GBM, HCT116, HELA, RPE1 | (Weng *et al.*, 2020) |
| CREB1 | Regulatory | {CREB1, ACACB} | DLD1, GBM, HCT116, HELA, RPE1 | (Fang *et al.*, 2016)<br>(Yang *et al.*, 2019) |
| CTPS2 | Metabolic | {CTPS2, TWIST1, TWIST2} | HELA | |
| PISD | Metabolic | {PISD, SPHK2, SPHK1, HLF} | HCT116 | |
| | | {PISD, SPHK2, BTG2, HLF} | HELA | (Bellance *et al.*, 2020) |
| RARA | Regulatory | {RARA, SCD5} | HELA | |
| STAT6 | Regulatory | {STAT6, CYP2R1, NFATC1, CYP27A1} | RPE1 | |
| TXN2 | Metabolic | {TXN2, PPARD} | DLD1 | Zhang et al., 2011 |
| | | | HELA | |
